# ANLN directly interacts with PCNA to regulate UV induced translesion synthesis

**DOI:** 10.1101/2024.08.27.609892

**Authors:** Bei-Bei Tong, Yu-Fei Cao, Bing Wen, Teng Fu, Dan-Xia Deng, Qian-Hui Yang, Yu-Qiu Wu, Hua-Yan Zou, Lian-Di Liao, Li-Yan Xu, En-Min Li

## Abstract

Anillin (ANLN) is a cytoskeletal binding protein involved in mitosis. ANLN is located in the nucleus during interphase and in the cytoplasmic contractile rings during mitosis. Our previous studies found that ANLN is abnormally overexpressed in esophageal squamous cell carcinoma (ESCC), promoting cell division by regulating contractile ring localization. However, the exact function of ANLN in the nucleus remains unclear. Here, we found that the expression of ANLN in the nucleus is associated with poor prognosis in ESCC patients, rather than in the cytoplasm. Protein mass spectrometry and bioinformatics analysis revealed that ANLN is related to DNA synthesis, and proliferating cell nuclear antigen (PCNA) is found to be a potential interacting protein of ANLN. PCNA directly interacts with the PIP box domain of ANLN and co-localizes in the nucleus. ANLN promotes DNA replication and S phase progression in a PCNA dependent manner and independent with the cytoskeletal function of ANLN. Importantly, ANLN is involved in transletion synthesis (TLS), a type of DNA synthesis under stress, by promoting PCNA monoubiquitination at K164 residue. Mechanistically, ANLN binds and recruits the E3 ligase RAD18 to promote PCNA monoubiquitination and DNA polymerase eta loading under UV radiation conditions. Consistently, depletion of ANLN leads to increased genomic instability and increased sensitivity to UV radiation. The findings of the study showed that ANLN in the nucleus as a protein scaffold is involved in UV induced DNA synthesis pathway, providing new insights into the function and mechanism of ANLN in cancer cells.

## Introduction

Anillin (ANLN) is a cytoskeletal protein involved in mitosis that promotes the localization of contractile rings and the process of cytokinesis by recruiting contractile ring components (1-4). ANLN contains 25 exons, encoding a protein with 1124 amino acid residues. The theoretical molecular weight of ANLN protein is 124 kDa, while SDS-PAGE electrophoresis shows a molecular weight of approximately 160 kDa (5). In 2015, Sun et al. identified the C-terminal domain (712-1124 aa) of ANLN protein through crystal structure analysis and found that it is related to plasma membrane binding. The N-terminal of ANLN contains F-actin and Myosin II binding domains, but the structure has not been resolved (6).

In previous reports, it was found that ANLN is highly expressed in various cancers and is associated with tumor malignant progression. Inhibiting the expression of ANLN can effectively inhibit the proliferation and invasion ability of cancer cells (7-11). Although most research focuses on the mitotic function of ANLN, in recent years, it has been found that ANLN may also function in the nucleus (8,12,13). ANLN depletion leads to transcriptional reprogramming and stemness suppression of breast cancer cells and the expression of ANLN in the nucleus, but not cytoplasmic ANLN, is negatively correlated with patient survival in urothelial carcinoma and renal cell carcinoma (12,14-16). These studies have found that nuclear ANLN may play a crucial role in cancer progression, but its exact function has not yet been identified. Our previous research has shown that high expression of ANLN promotes contractile ring localization, cytokinesis, and proliferation in esophageal squamous cell carcinoma (ESCC) cells. Subsequently, we further discovered that DNA synthesis related proteins are associated with ANLN, such as PCNA. Therefore, we consider that nuclear ANLN may be related to the process of DNA synthesis, which may be an important factor for ANLN to promote ESCC cell proliferation (5).

PCNA is a core functional protein of DNA synthesis (17). It is a conserved protein expressed in most eukaryotic cells, typically forming homotrimer rings that can contain and slide along double-stranded DNA (dsDNA) (18,19). In the S phase, PCNA is loaded at the primer-template junction with help of the Replication Factor C (RFC) complex, then interacts with the high-fidelity replicative DNA Polymerases ε and δ to promote DNA replication (20,21). When cells experience replication stress, DNA lesions would block the progression of high-fidelity DNA Polymerases. In response to this situation, DNA damage tolerance pathways are evolved for the cellular survival. Translesion DNA synthesis (TLS) is an important part of DNA damage tolerance (22-24). It bypass DNA damage site with specific low-fidelity polymerases such as Pol ι, Pol η, Pol κ and Rev1. These TLS polymerases belong to the Y-family and the high-fidelity DNA polymerases Pol α, Pol δ, and Pol ε, which belong to the classical β-family. PCNA plays a key role in DNA polymerases exchange (25-27). In the process of replication stress triggering TLS, RPA binds to ssDNA and recruits the E3 ubiquitin ligase RAD18 (28). K164 residue of PCNA was monoubiquitinated by RAD18 to enhance its binding capacity with TLS polymerases (29).

In this study, it was found that ANLN interacts with PCNA in vitro and vivo. We also found that ANLN promotes DNA synthesis and cell proliferation in a PCNA dependent manner. Additionally, under UV induced replication stress, ANLN promotes the PCNA monoubiquitin and TLS pathways by recruiting RAD18. Consistent with this, ANLN is crucial for the DNA synthesis and UV radiation sensitivity of ESCC cells.

## Results

### Nuclear ANLN is associated with DNA replication proteins in ESCC cells

To reveal the role of ANLN in ESCC cells, immunofluorescence analysis was used to investigate the subcellular localization of ANLN. The results showed that ANLN was located in the nucleus during the interphase and in the contractile rings during mitosis (Fig. 1A). Consistently, cellular fractionation showed the presence of ANLN in cytoplasm and nucleoplasm and on chromatin (Fig. 1B). To reveal the mechanism of ANLN in the nucleus promoting malignant proliferation of ESCC cells, gene set enrichment analysis (GSEA) was performed on the proteomics of 124 ESCC patients (IPX0002501000) (30). Results showed that the expression level of ANLN protein was positively correlated with DNA replication (Fig. 1C). To confirm this correlation, ANLN knockdown cells were detected using EdU incorporation experiments. Downregulation of ANLN expression leads to a decrease in the proportion of cells undergoing DNA replication (Fig. 1D-F). We conclude that nuclear ANLN is associated with DNA replication in ESCC cells.

**Figure 1.**
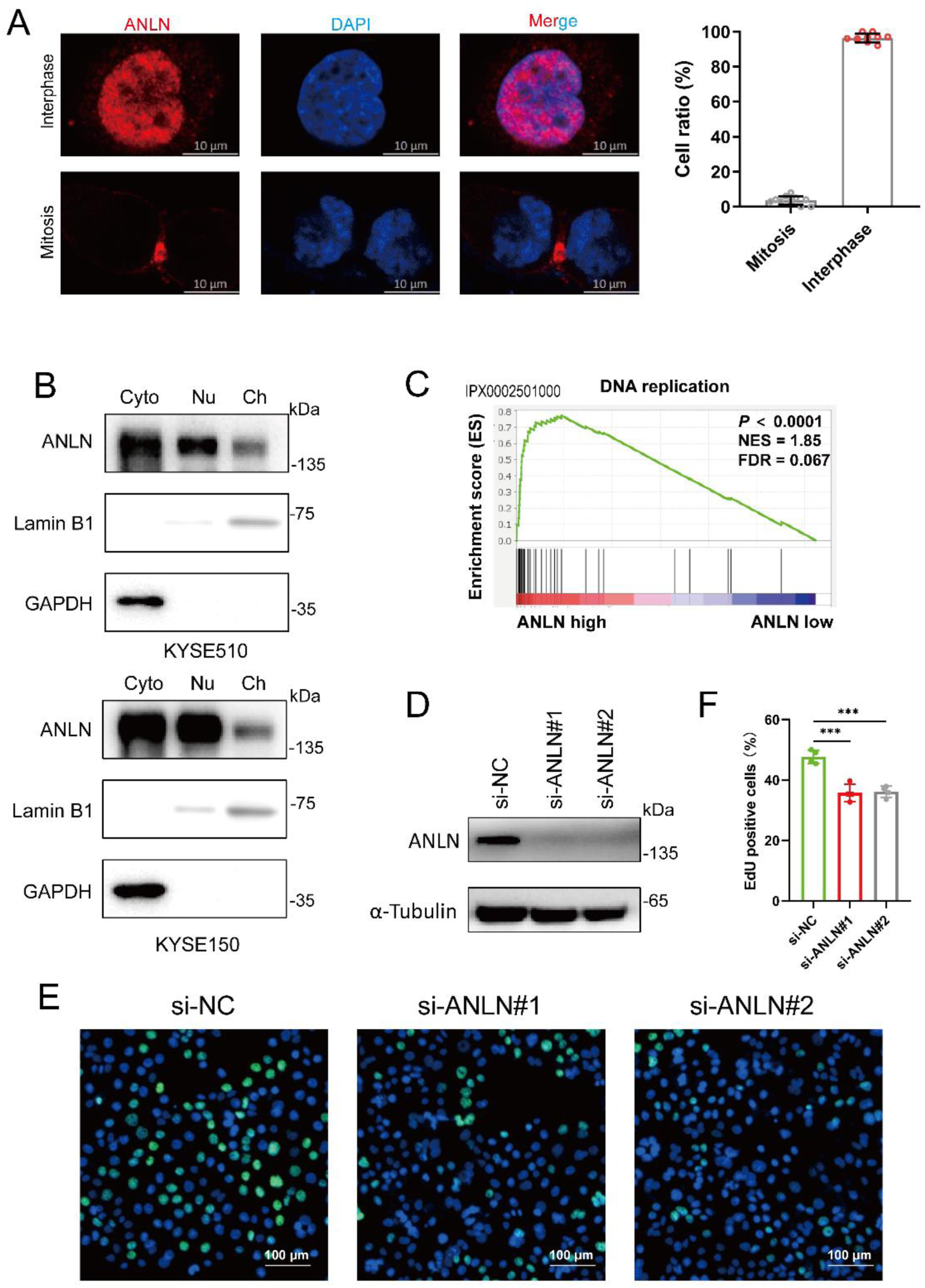
Nuclear ANLN is associated with DNA replication proteins in ESCC cells. (**A**) Representative immunofluorescence image of ANLN subcellular localization in KYSE150 cells during mitosis and interphase (Left). The cell ratio of different staining pattern (Right). (**B**) Western blotting was performed to examine the different subcellular fractions of ESCC cells. GAPDH was used as a cytoplasmic marker and Lamin B1 was used as a chromatin marker. Cyto, cytoplasm; Nu, nucleus; Ch, chromatin. (**C**) Based on the GSEA of 124 ESCC tissues, analyze the potential function of ANLN in ESCC. (**D-F**) KYSE150 cells were transfected with siRNA for 48 h, and the proportion of S phase was detected by EdU assay. Representative images determined by EdU (E). The statistical results of EdU assay (three parallel data, mean ± S.D, unpaired *t*-test, \*\*\**P* < 0.001) (F). Western blotting was used to detect the knockdown effect of ANLN (D).

### ANLN regulates S phase progression of ESCC cells in an Actin-independent manner

To explore whether ANLN promotes DNA replication and S phase progression, cell cycle synchronization model was constructed using thymidine to validate the effect of ANLN on the S phase progression (Fig. 2A). Immunoblotting showed that cyclin E2 was highly expressed at the G1/S phase boundary and degraded upon entering the S phase (31). However, depletion of ANLN resulted in delayed degradation of cyclin E2, indicating a slowdown in the S phase process (Fig. 2B). Consistently, flow cytometry also showed that inhibiting ANLN expression in ESCC cells resulted in delayed S phase progression (Fig. 2C). To investigate the effect of ANLN expression on DNA replication, ANLN knockdown cells were detected using DNA fiber assay. The cells were labeled with iododeoxyuridine (IdU) for 30 min, and then chlorodeoxyuridine (CldU) for 30 min. DNA fibers were observed by immunofluorescence (Fig. 2D). As expected, ANLN depletion decreased DNA fiber tract length and DNA replication initiation in ESCC cells (Fig. 2E, F). It is interesting that ANLN knockdown also inhibited DNA replication in normal human cells (Supplementary Fig. 1A-D).

**Figure 2.**
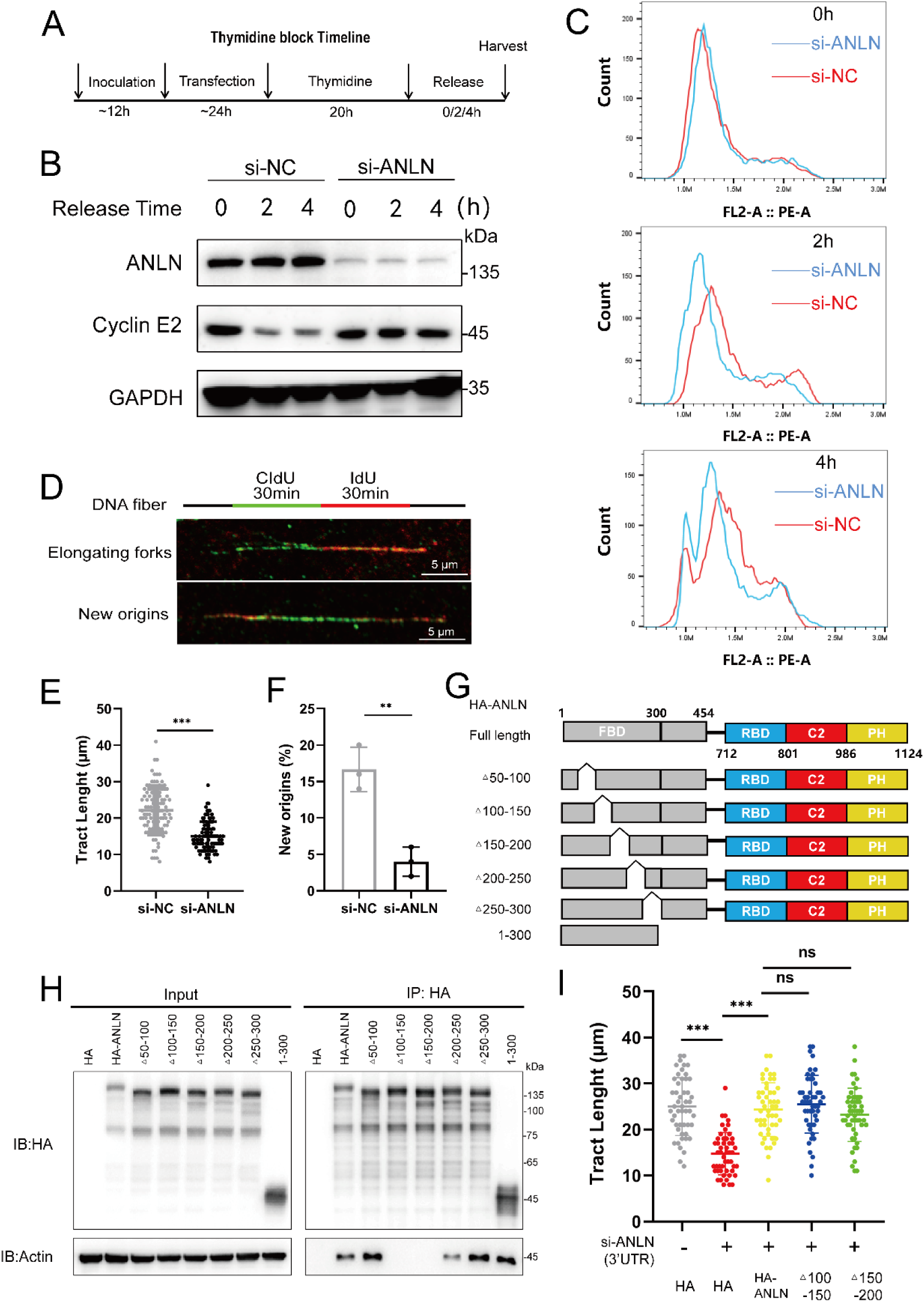
ANLN regulates S phase progression of ESCC cells in an Actin-independent manner. (**A**) Flow chart of cell cycle synchronization. (**B**) Expression of ANLN and cyclin E2 was detected by western blotting. (**C**)The effect of ANLN knockdown on the S phase progression of KYSE150 cells was measured by flow cytometry. (**D-F**) Schematic of DNA fiber assay, after 48 h of transfection, the cells were labeled with 50 μM CldU and 250 μM IdU for 30 min each. Representative images of elongating fork and new origin (D). Statistical analysis of the tract length (E) (100 fibers were analyzed per condition, mean ± S.D, unpaired *t*-test, \*\*\**P* < 0.001) and new origin of DNA fiber (F) (three parallel data, 50 fibers in each group, mean ± S.D, unpaired *t*-test, \*\*\**P* < 0.001). (**G**) Schematic illustration of ANLN structure. (**H**) The indicated plasmids were transfected into HEK293T cells, the interaction of ANLN mutants and PCNA was detected by co-immunoprecipitation. (**I**) Tract length analysis of DNA fiber assay, the indicated plasmids were stable overexpression in KYSE150 cells, after 48 h of siRNA (Targeting 3 ’UTR) transfection, the cells were labeled with 50 μM CldU and 250 μM IdU for 30 min each (50 fibers were analyzed per condition, mean ± S.D, unpaired *t*-test, \*\*\**P* < 0.001).

Next, we asked if ANLN regulates DNA replication by its cytoskeletal function. In previous reports, ANLN assists in mitosis by interacting with Actin (1,6). Here, we validated the specific domain of the ANLN-Actin interaction and found that ANLN did not interact with Actin after the absence of amino acid residues at positions 100-150 or 150-200 aa (Fig. 2G, H). Then, we knocked down the expression of endogenous ANLN in KYSE150 cells and then rescue with ANLN mutants. DNA fiber assay showed that the actin-binding deficient ANLN mutant was equally active as wild type in recover DNA replication (Fig. 2I). Therefore, we believe that the effect of ANLN on DNA replication is independent of its cytoskeleton-related functions.

### ANLN directly interacts with PCNA

To test that how ANLN participate in DNAreplication, the interaction protein mass spectrometry of ANLN was analyzed for exploring which DNA replication functional proteins interact with ANLN (5). At first, we screened 2063 chromatin binding-related proteins from UniProt database. Among them, 265 proteins were screened by ANLN interaction proteomics and ESCC proteomics (Supplementary Fig. 2A-B). And within that, 15 proteins were overexpressed in ESCC proteomic data (Supplementary Fig. 2C). 6 of 15 proteins is involved in DNA replication. Among them, only PCNA was an independent prognostic poor factor for patients with ESCC (Supplementary Fig. 2D). PCNA is a classic DNA synthesis related protein that is highly expressed in ESCC tissues and closely associated with poor prognosis in patients (30,32) (Supplementary Fig. 2E, F). Therefore, we propose that PCNA is the key interaction protein in which ANLN participates in DNAreplication and affects the malignant proliferation of ESCC.

To confirm whether there is an interaction between ANLN and PCNA. HA-ANLN and Flag-PCNA plasmids were next transfected into HEK293T cells then subjected to immunoprecipitation experiments. The results suggest that ANLN and PCNA are in the same protein complex and may interact directly with each other (Fig. 3A). Consistent with this, experimental results in ESCC cells showed that endogenous ANLN and PCNA are also in the same protein complex (Fig. 3B). Furthermore, the interaction was detected using a proximity alignment assay (PLA) and found that it was localized in the nucleus (Fig. 3C, D). It has been found that PCNA interacting proteins typically contain a conserved PCNA-interacting proteins box (PIP-box) sequence (33,34). Using a sequence alignment, we found a PIP box (405-411 KAIQERL) in the human ANLN protein (Fig. 3E). Therefore, the 405-411 amino acid region of ANLN was deleted to construct a mutant plasmid, and then co-transfected with Flag-PCNA into HEK293T cells for immunoprecipitation experiments. As expected, the deletion of the 405-411 amino acid region disrupted the interaction between ANLN and PCNA (Fig. 3F). Consistently, PCNA interacts with the N-terminal 1-454 region of ANLN containing the PIP box, rather than the C-terminal 454-1124 region (Fig. 3F). Expression and purification of His labeled ANLN and PCNA without label proteins from prokaryotic cells, followed by in vitro His pulldown assay. Consistent with the result in vivo, ANLN directly interacts with PCNA through the PIP box (Fig. 3G). In addition, we notice that PCNA forms foci at sites of origin firing and DNA synthesis in previous studies (35). We simultaneously imaged GFP-ANLN and RFP–PCNA chromobody (PCNA-CB) in different cell cycles by living cell microscope (36). The results showed that ANLN not only dispersed in the nucleus, but also always existed at the foci of PCNA during the S phase (Fig. 3H). In the mitotic stage, consistent with previous conclusions, there was no obvious co-localization of ANLN and PCNA (Fig. 3H). In summary, ANLN interacts directly with PCNA in the nucleus through PIP box.

**Figure 3.**
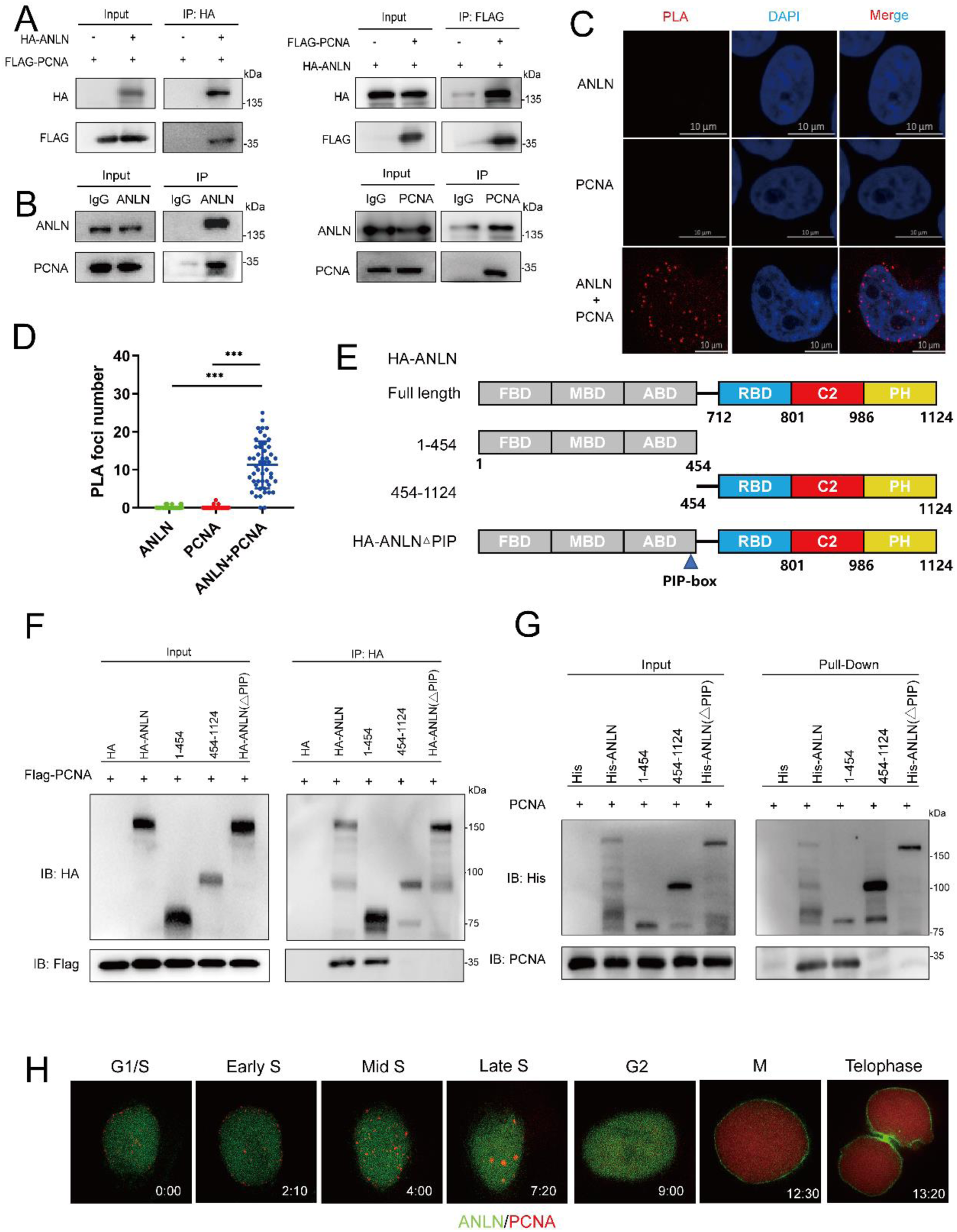
ANLN directly interacts with PCNA. **(A)** The indicated plasmids were transfected into HEK293T cells, and the interaction between exogenous ANLN and PCNA was detected by co-immunoprecipitation. (**B**) The interaction between endogenous ANLN and PCNA was detected by immunoprecipitation in KYSE150 cells. (**C**) The interaction between ANLN and PCNA was detected by PLA assay. (**D**) PLA foci numbers of 50 cells in each group were analyzed (three parallel data, mean ± S.D, unpaired *t*-test, \*\*\**P* < 0.001). (**E**) Schematic illustration of ANLN structure. (**F**) The indicated plasmids were transfected into HEK293T cells, the interaction of ANLN mutants and PCNA was detected by co-immunoprecipitation. (**G**) The interaction between PCNA and different domains of His-ANLN was detected by His pulldown assay. (**H**) Representative images of ANLN and PCNA during interphase and mitosis in KYSE150 cells.

### ANLN regulates S phase progression of ESCC cells in a PCNA dependent manner

To determine whether ANLN promotes DNA replication through PCNA dependent manner, ANLN wild-type or PIP box deletion mutants were reintroduced into Dox induced ANLN knockdown cells (Fig. 3A). The EdU incorporation experiment showed that reintroducing ANLN wild-type instead of PIP-box deletion mutant rescued ANLN knockdown induced S phase cell reduction (Fig. 4A, D). Similarly, colony formation experiments showed that ANLN promotes ESCC cell proliferation through the interaction with PCNA. (Fig. 4B, E). Moreover, DNA fibers assay proved that ANLN-△PIP could not restore normal DNA replication rate (Fig. 4F). It is suggested that ANLN is involved in DNA replication in PCNA dependent manner. We next investigated whether ANLN regulates the localization of PCNA (37). PCNA is loaded at the primer-template junction in S phase, thus we obtained G1/S cells by thymidine block. Cell fractionation revealed that ANLN knockdown reduced the level of PCNA in chromatin components, suggesting that ANLN is associated with PCNA chromatin loading (Fig. 4G).

**Figure 4.**
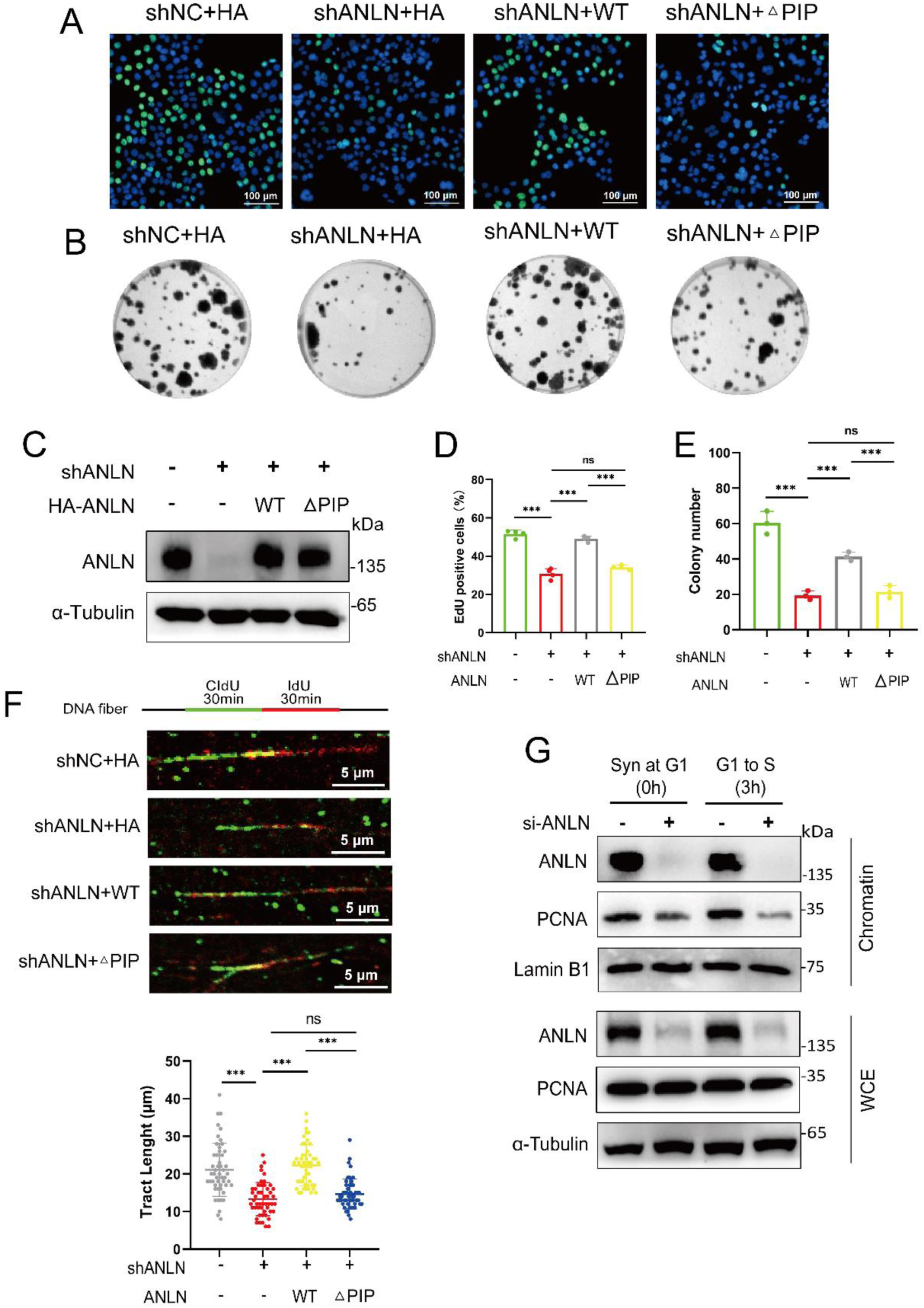
ANLN regulates S phase progression of ESCC cells in a PCNA dependent manner. (**A-F**) Dox (2 μg/mL) induced ANLN knockdown in KYSE150 cells and transfected with the indicated plasmids; ANLN expression was analyzed by western blotting (C). (**A, D**) EdU assay measured the S phase proportion of KYSE150 cells. Representative image of EdU (three parallel data, mean ± S.D, unpaired *t*-test, \*\*\**P* < 0.001). (**B, E**) The proliferation of ESCC cells was analyzed by colony formation assay (three parallel data, mean ± S.D, unpaired *t*-test, \*\*\**P* < 0.001). (**F**) Schematic of DNA fiber assay (Up). Statistical analysis of the tract length (Below) (50 fibers were analyzed per condition, mean ± S.D, unpaired *t*-test, \*\*\**P* < 0.001). (**G**) ANLN knockdown KYSE150 cells were synchronized in late G1 by thymidine block followed or not with 3 h release into S phase. PCNA chromatin binding was detected by cellular fractionation. α-Tubulin was used as a cytoplasmic marker and Lamin B1 was used as a chromatin marker. WEC, whole-cell lysate.

Previous studies have reported that abnormal mitosis can cause DNA damage and inhibit replication (38). To exclude the interference of mitotic defects caused by ANLN mutations, the cell model was detected through synchronization and immunofluorescence. Results showed that the staining pattern of ANLN wild-type and △PIP were consistent, both in the nucleus and contractile ring. In addition, PIP-box deletion did not cause furrow localization defect of the contractile ring (Supplementary Fig. 3A-D). Taken together, these results indicate that ANLN regulates PCNA chromatin localization and DNA replication, thereby promoting S phase progression and proliferation of ESCC cells.

### ANLN participates in UV induced translesion synthesis by promoting PCNA monoubiquitination

Given that ANLN plays an important function in normal DNA replication, we next investigated whether it also plays a role during replication stress. DNA damage derived from endogenous and exogenous sources often occurs in the process of cell proliferation (27,39,40). When cells experience replication stress, the E3 ubiquitin ligase RAD18 mediates PCNA (K164) monoubiquitination, which induces low-fidelity and low-speed TLS polymerases combining with PCNA to complete translesion synthesis (TLS). These TLS polymerases belong to the Y-family include Pol ι, Pol η, Pol κ and Rev1. Among them, DNA Pol η (Pol eta) can correctly bypass UV induced cyclobutane pyrimidine dimers (CPDs) damage (41-43). Activated Pol eta avoids DNA replication arrest, replication fork collapse, or double strand breaks (DSBs), thereby the forks can continue to replicate across the DNA damage site.

To investigate the effect of ANLN on the TLS pathway, an ESCC cell model simulating replication stress was constructed using UV irradiation. ESCC cells were treated with different doses of UV, and then cultured for 2 hours. The UV irradiation dose was determined based on the level of PCNA monoubiquitination. It was found that 100J/m^2^ UV treatment significantly enhanced the monoubiquitination level of PCNA (Supplementary Fig. 4A), while ANLN depletion prevented UV induced monoubiquitination of PCNA but did not affect the protein expression level of PCNA (Fig. 5A and Supplementary Fig. 4B).

**Figure 5.**
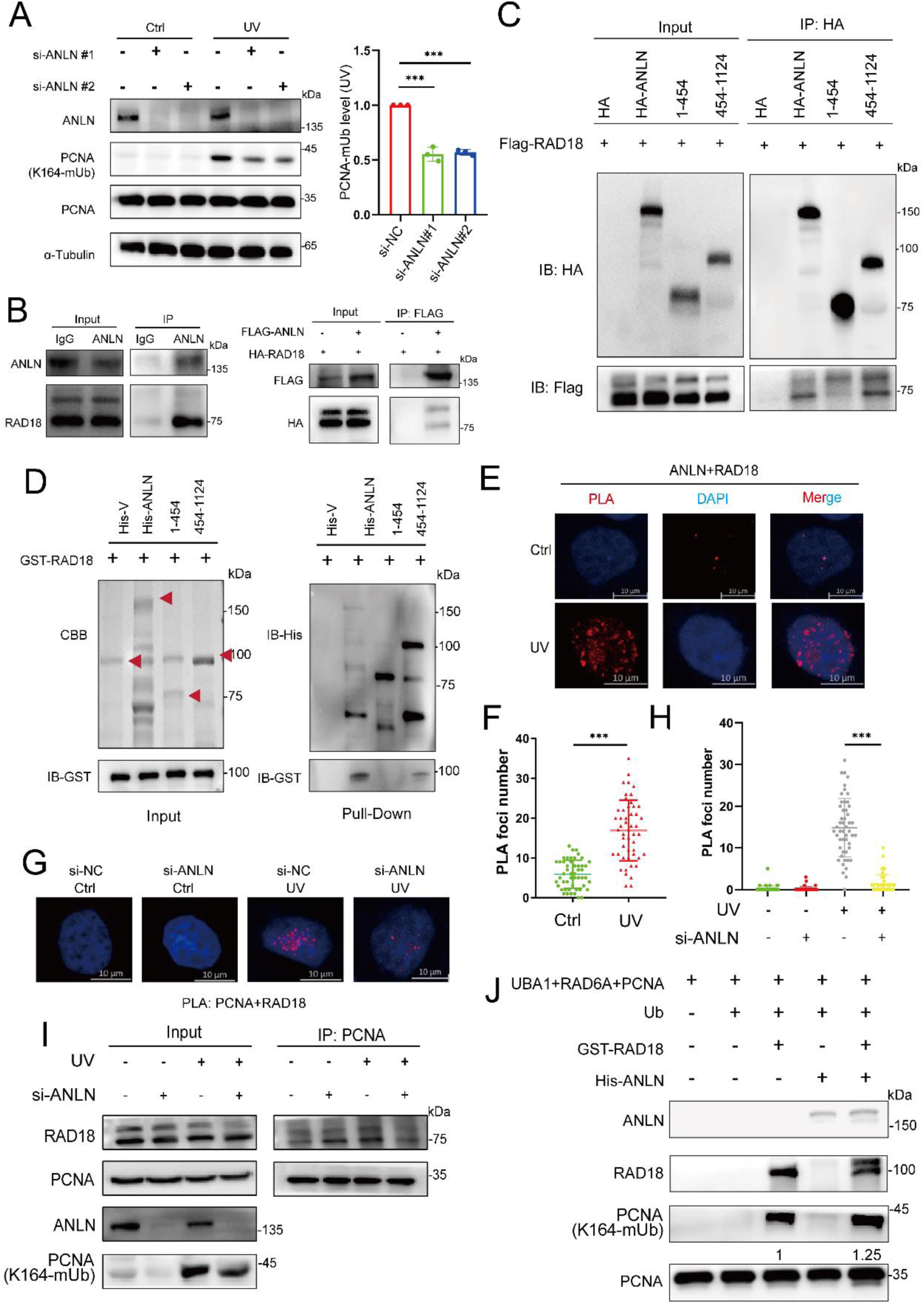
ANLN participates in UV induced translesion synthesis by promoting PCNA monoubiquitination. (**A**) KYSE150 cells transfected with siRNAs were treated with UV. The monoubiquitination level of PCNA was measured by western blotting (Left). Statistical results of three independent repeated experiments (Right) (three parallel data, mean ± S.D, unpaired *t*-test, \*\*\**P* < 0.001). (**B**) The interaction between endogenous ANLN and RAD18 was detected by immunoprecipitation in KYSE150 cells (Left). The indicated plasmids were transfected into HEK293T cells, ANLN and RAD18 binding were detected by immunoprecipitation (Right). (**C**) The indicated plasmids were transfected into HEK293T cells, the interaction of ANLN mutants and RAD18 was detected by co-immunoprecipitation. (**D**) The interaction between RAD18 and different domains of His-ANLN was detected by His pulldown assay. (**E, F**) After UV irradiation, the interaction between ANLN and RAD18 in KYSE510 cells was analyzed by PLA assay (E). PLA foci numbers of 50 cells in each group were analyzed (F). (three parallel data, mean ± S.D, unpaired *t*-test, \*\*\**P* < 0.001). (**G, H**) KYSE510 transfected with siRNAs was treated with UV, the interaction between PCNA and RAD18 was analyzed by PLA assay (G). PLA foci numbers of 50 cells in each group were analyzed (H) (three parallel data, mean ± S.D, unpaired *t*-test, \*\*\**P* < 0.001). (**I**) KYSE150 transfected with siRNAs was treated with UV, the interaction between PCNA and RAD18 was detected by immunoprecipitation. (**J**) Monoubiquitination of PCNA. Flag-PCNA was purified from HEK293T cells and monoubiquitinated by adding UBE1, RAD6B, RAD18, ANLN and Ub.

Since ANLN is usually functions as a protein scaffold to mediate protein recruitment, we next asked whether ANLN promotes PCNA monoubiquitination by recruiting RAD18. The correlation between ANLN and RAD18 was revealed by protein-protein interaction methods. Immunoprecipitation showed that ANLN interacts with RAD18 in ESCC or HEK293T cells (Fig. 5B). Next, ANLN and its mutant plasmids were transfected into HEK293T cells and analyzed using immunoprecipitation. Results showed that RAD18 interacts with the C-terminal 454-1124 amino acid region of ANLN (Fig. 5C). GST-RAD18 and His-ANLN mutant proteins were expressed in prokaryotic cells and purified for His pulldown analysis. Consistently, the RAD18 purified protein directly interacts with the C-terminus of ANLN (454-1124) instead of the N-terminus (1-454) (Fig. 5D). Interestingly, the interaction between ANLN and RAD18 was significantly enhanced after UV treatment, suggesting the potential role of ANLN in recruiting RAD18 (Fig. 5E, F). To reveal whether ANLN promotes the recruitment of RAD18 under replication stress, the interaction between RAD18 and PCNA in ESCC cells was analyzed after UV treatment. Both PLA and immunoprecipitation experiments showed that the enhanced interaction between RAD18 and PCNA induced by UV radiation was reversed by ANLN knockdown (Fig. 5G-I). In addition, we performed PCNA ubiquitination in vitro. It is found that adding ANLN could increase the ubiquitin modification level of PCNA by RAD18 to a certain extent (Fig. 5J). Overall, these results indicate that ANLN promotes PCNA monoubiquitination and participates in the TLS pathway by recruiting RAD18.

### ANLN regulates tolerance to genomic DNA damage under DNA replication stress

Pol eta plays a crucial role in preventing tumor cell death during replication stress recovery, bypassing UV induced CPD, and avoiding genomic instability after UV induced DNA damage (44). Monoubiquitinated PCNA promotes the TLS pathway by recruiting Pol eta, while knockdown of ANLN leads to decreased PCNA ubiquitination (Fig. 5A). Therefore, we next investigated whether ANLN affects UV induced Pol eta chromatin loading. Cellular fractionation revealed that UV induced increase in Pol eta chromatin binding was reduced by ANLN knockdown (Fig. 6A). Immunoprecipitation and PLA assay also revealed that knockdown of ANLN caused a decrease in the binding between PCNA and Pol eta under UV conditions (Fig. 6B-D). Furthermore, an alkaline comet assay was performed to detect DSB levels in ESCC cells. The results showed that inhibiting the expression of ANLN significantly enhanced the level of DSB induced by UV, suggesting an increase in genomic instability (Fig. 6E, F). We then used clone formation experiments to test the survival rate of ESCC cells after UV irradiation. The results indicate that ANLN depletion leads to increased sensitivity of cells to UV radiation, while reintroduction of HA-ANLN restores cell resistance to UV radiation (Fig. 6G-H). Collectively, the expression of ANLN is negatively correlated with UV sensitivity in ESCC cells.

**Figure 6.**
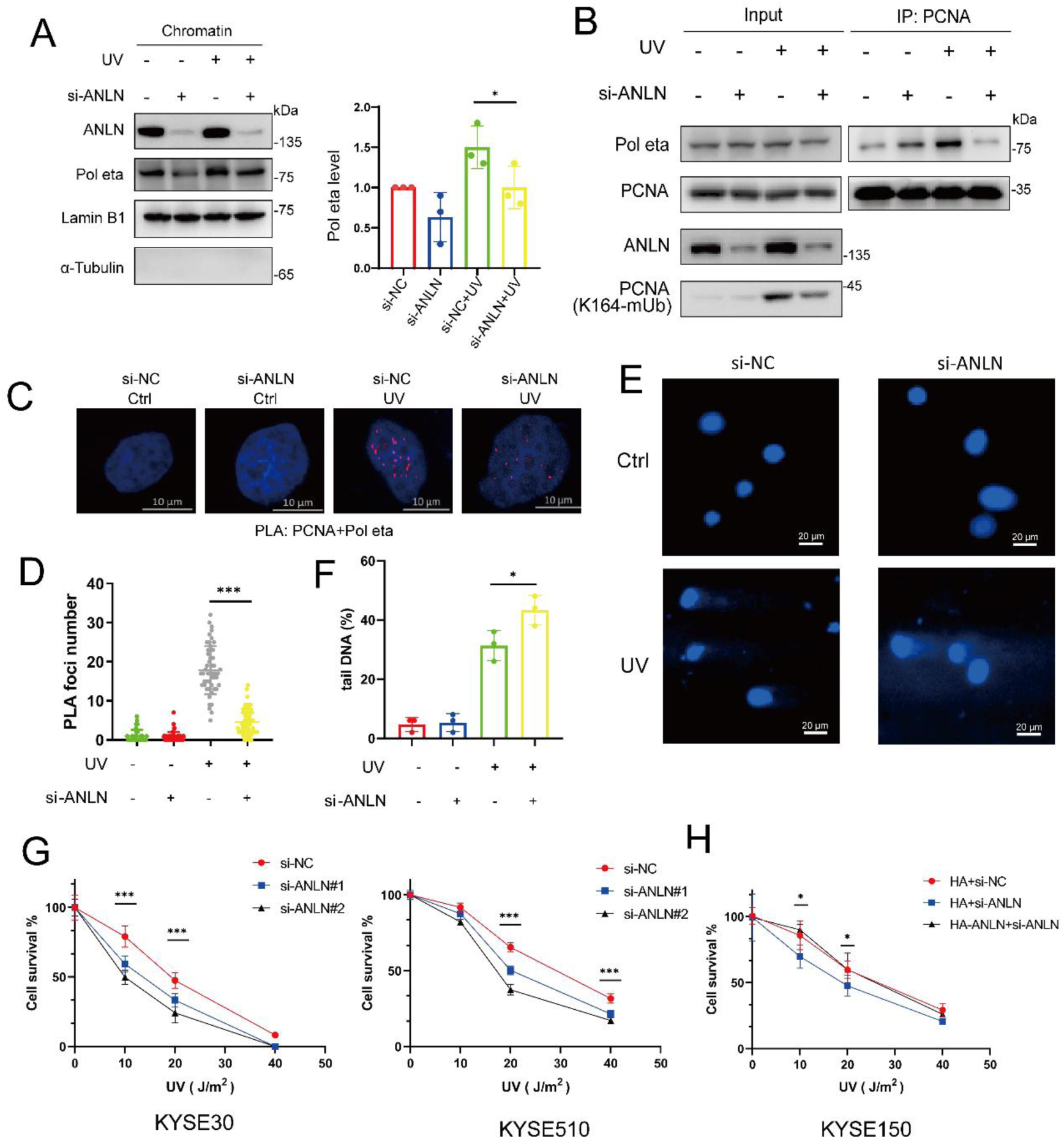
ANLN regulates tolerance to genomic DNA damage under DNA replication stress. (**A**) KYSE150 transfected with siRNAs was treated with UV, the chromatin binding of Pol eta was detected by cellular fractionation. α-Tubulin was used as a cytoplasmic marker, and Lamin B1 was used as a chromatin marker (Left). Statistical results of three independent repeated experiments (Right) (three parallel data, mean ± S.D, unpaired *t*-test, \*\*\**P* < 0.001) (**B**) KYSE150 transfected with siRNAs was treated with UV, the interaction between PCNA and Pol eta was detected by immunoprecipitation. (**C, D**) KYSE510 transfected with siRNAs was treated with UV, the interaction between PCNA and Pol eta was analyzed by PLA assay (C). PLA foci numbers of 50 cells in each group were analyzed (D). (three parallel data, mean ± S.D, unpaired *t*-test, \*\*\**P* < 0.001). (**E, F**) KYSE150 transfected with siRNAs was treated with UV, the level of DNA damage was measured by an alkaline comet assay (E). The percentage of DNA in the comet tail, with 50 cells counted for each group (F). (three parallel data, mean, ± S.D, unpaired *t*-test, \**P* < 0.05). **(G, H)** ESCC cells transfected with siRNAs or stably expressing indicated plasmids were treated with UV, and then cell proliferation was detected using a colony formation assay. (three parallel data, mean ± S.D, unpaired *t*-test, \**P* < 0.05, \*\*\**P* < 0.001).

### High expression of ANLN in the nucleus is closely related to the poor prognosis of ESCC patients

We previously analyzed tissue chips from 104 ESCC patients and found that the expression of ANLN protein in ESCC tissues was significantly higher than that in normal tissues (5). To demonstrate the clinical significance of nuclear ANLN, further analysis was carried out on immunohistochemical data from previous studies. The results showed that ANLN was expressed in the nucleus of both normal and ESCC tissues (Fig. 7A and Supplementary Fig. 5). We found that high expression of ANLN in the nucleus was negatively correlated with overall survival in patients, but not with ANLN in the cytoplasm (Fig. 7B, C). The cox regression analysis showed that the high expression of ANLN in the nucleus was not affected by age, gender, smoking, drinking and other factors of patients with ESCC, and was an independent prognostic factor for patients with ESCC. Although the cytoplasm ANLN showed a similar effect, it was not statistically significant (Fig. 7D, E). These results suggest that abnormally high expression of ANLN in the nucleus promotes the malignant proliferation of ESCC cells and is closely associated with the poor prognosis of patients with ESCC.

**Figure 7.**
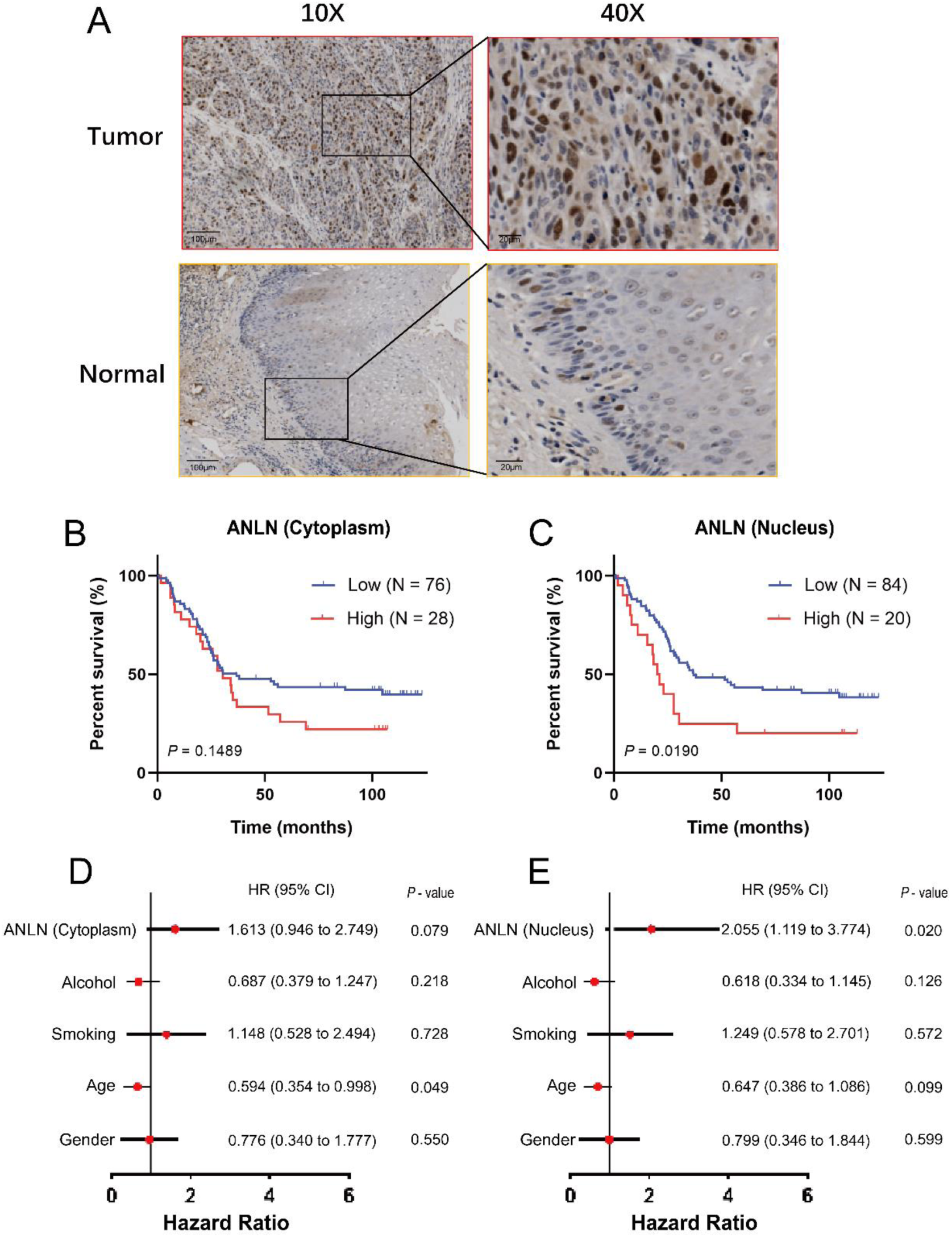
High expression of ANLN in the nucleus is closely related to the poor prognosis of ESCC patients. (**A**) Representative image of ANLN immunohistochemical staining in ESCC and normal tissues (scale 100 and 20 μm). (**B, C**) Kaplan–Meier curve analysis of the correlation between nuclear or cytoplasmic ANLN protein expression and overall survival of 104 ESCC patients. (**D, E**) Forest plot of the result of Cox proportional hazards regression analysis of ANLN and clinical information of 104 ESCC patients.

## Discussion

ANLN promotes malignant progression of cancer cells by mediating the contractile ring organization and cytokinesis (7-9,12). However, whether ANLN in the nucleus contributes to cancer progression remains unclear. Here, we report that the nuclear ANLN directly interacts with PCNA, promoting DNA replication and ESCC cell proliferation. Surprisingly, under UV induced replication stress, ANLN promotes PCNA monoubiquitination by recruiting RAD18, thereby promoting Pol eta loading and activation of the TLS pathway. Moreover, ANLN is negatively correlated with genomic stability and sensitivity to UV radiation, suggesting that targeting ANLN can reduce the DNA damage tolerance of ESCC cells. Finally, it was found that high expression of ANLN in the nucleus is associated with poor prognosis in ESCC patients. These results provide new insights into the function and mechanism of ANLN in cancer, and establish a novel theoretical basis for intervention strategies in ESCC.

The loading of PCNA onto chromatin is mediated by ATP-dependent clamp loader replication factor C (RFC) (45). We demonstrate that ANLN promotes the chromatin loading of PCNA (Fig. 3J), suggesting that ANLN may cooperate with the RFC complex to mediate PCNA localization. Consistent with this, several RFC complex subunits were identified by ANLN interacting protein mass spectrometry. Therefore, ANLN may participate in DNA replication by organizing the RFC complex and loading PCNA on chromatin, which requires further research in the future.

Various types of DNA damage often occur during the proliferation process of cancer cells, and the TLS pathway avoids replication arrest or replication fork collapse induced by DNA damage (27). In the TLS pathway, PCNA recruits low-fidelity DNA polymers (Pol eta) at damaged DNA sites to replace high-fidelity DNA polymers (Pol delta), resuming replication and avoiding replication fork collapse. We showed that after UV treatment of ESCC cells, ANLN enhanced the interaction between RAD18 and PCNA, promoting chromatin loading of Pol eta. Additionally, ANLN knockdown leads to enhanced sensitivity of ESCC cells to UV irradiation. Whether this is mediated by TLS pathway alone needs further exploration. Protein interaction mass spectrometry showed that ANLN is also associated with several DNA damage repair proteins, including nucleotide excision repair and mismatch repair.

Overall, this study suggests that nuclear ANLN promotes DNA replication and proliferation by mediating PCNA chromatin loading, and also participates in the TLS pathway, promoting proliferation by enhancing DNA damage tolerance.

## Experimental procedures

### Data Sources and Bioinformatic Analysis

The immunohistochemical data of ANLN in ESCC tissues and the mass spectrometry data of ANLN interacting proteins were obtained from our published paper (5,30). The proteomic data of 124 ESCC tissues were obtained from the iProX database (IPX0002501000) (30). Chromatin binding related proteins were downloaded from the UniProt database (https://www.uniprot.org/) and then overlapped with ANLN interacting proteins to obtain candidate proteins for this study. The candidate proteins were analyzed in the Metscape database (https://metascape.org/), and an enrichment analysis was performed using KEGG (46). SPSS software was used to calculate the differential expression analysis (Wilcoxon matched pairs signed rank test) and survival analysis (Kaplan-Meier analysis) of these proteins. While a two-tailed p-value less than 0.05 was considered to indicate statistical significance.

### Cell culture

ESCC cell lines KYSE30, KYSE150 and KYSE510 were cultured in RPMI-1640 medium (GIBCO) supplemented with 10% fetal bovine serum (GIBCO), 100 U/mL penicillin G, and streptomycin. HEK293T cell lines were cultured in Dulbecco’s modified Eagle’s medium (GIBCO), with 10% fetal bovine serum. Cells were incubated in humidified atmosphere at 37℃ and 5% CO_2_. Cell lines were authenticated by short tandem repeat profiling and were routinely tested for mycoplasma contamination (47,48).

### Immunofluorescence assay

Cells were fixed with 4% paraformaldehyde for 15 min, permeabilized with 0.1% Triton X-100 for 10 min, then the cells were blocked with 5% bovine serum albumin for 1 h. After that, cells were incubated overnight with antibodies against ANLN (1:1000, Abcam, ab211872) and PCNA (1:200, Proteintech, 10205-2-AP). Moreover, cells were incubated with Alexa fluor 647 donkey anti-mouse IgG (Jackson, 715-605-150, 1:200), Alexa fluor 488 donkey anti-rabbit IgG (Jackson, 711-545-152, 1:200) in the dark for 1 h. Finally, cells were stained with DAPI (Beyotime, C1005, 1:2000) in the dark for 10 min and observed by LSM800 with airyscan confocal laser scanning microscope (Carl Zeiss, Oberkochen, Germany) (5,49).

### Cell Fractionation

Cells were cultured in 10 cm dishes and digested using trypsin, collected by centrifugation at 600× g for 3 min, washed with PBS, and lysed with 600 µL of hypotonic buffer (10 mM Tris-HCl, PH 7.5, 2 mM MgCl_2_, 3 mM CaCl_2_, 320 mM sucrose, 1 mM dithiothreitol (DTT), 1% protease inhibitor (MedChemExpress, HY-K0010) and 0.3% NP40) on ice for 10 min. Next, the cells were centrifuged for 5 min at 2800× g, and the supernatant contained cytoplasmic proteins. The pellet was washed with hypotonic buffer for three times, and then incubated with 100 µL nuclear extraction buffer (20 mM HEPES, pH 7.7, 420 mM NaCl, 1.5 mM MgCl_2_, 200 mM ethylene diamine tetraacetic acid (EDTA), 25% glycerol, 1% protease inhibitor and 1 mM DTT) on ice for 30 min, and then centrifuged at 8000× g for 15 min. The supernatant contained soluble nuclear proteins and the chromatin pellet was lysed with 1×SDS loading buffer, sonicated, and analysised with western blotting (50).

### Western blotting

Western blotting performed as previously described (5). Cells were lysed in the SDS sample loading buffer, samples were boiled for 15 min and loaded on 8% SDS-PAGE. The proteins were transferred onto polyvinylidenefluoride (PVDF) membranes, and then blocked with 5% skim milk for 1 h. Membranes were incubated with antibodies against ANLN (1:2000, Abcam, ab99352), GAPDH (1:3000, Proteintech, 60004-1-Ig), PCNA (1:2000, Proteintech, 10205-2-AP), HA-tag rabbit antibody (1:3000, Cell Signaling Technology, C29F4), Flag M2 monoclonal antibody (1:5000, Sigma, F3165-1MG), His-tag mouse antibody (1:1000, TransGen Biotech, HT501-01), cyclin E2 (1:1000, Cell Signaling Technology, 4132), α-Tubulin (1:10000, Sigma, T9026), PCNA-K164-Ub (1:1000, CST, 13439S), Pol eta (1:1000, Proteintech, 28133-1-AP), RAD18 (1:1000, Proteintech, 18333-1-AP). Proteins were detected by incubation with horseradish peroxidase-conjugated secondary antibodies and the signals were enhanced by luminol reagent (Bio-sharp, BL520B). Finally, protein signals were detected using ChemiDoc Touch (Bio Rad) and grayscale values were analyzed using Image Lab software version 2.0 (51).

### Small interfering RNA (siRNA) knockdown

Experiments were performed according to the manufacturer’s instructions. Single siRNA oligonucleotides targeting human ANLN and negative control siRNA were diluted by Opti-MEM® I Reduced Serum Medium (Life Technology, 31985-070) and then mixed with siRNA Transfection Reagent (Life Technology, 13778150). Fresh medium was replaced 8 h after transfection, and the cells were further cultured for 40 h, followed by subsequent experiments (5). The siRNAs targeted ANLN (siANLN#1: 5’-CCA GAC CUC UGC UUU CAA ATT-3’, and siANLN#2: 5’-GCA GAU ACC AUC AGU GAU UTT-3’) were synthesized by GenePharma.

### Lentiviral production and infection

The shRNA targeted ANLN (5’-CCG GGC CAA TAT TCA CTA CGT ATT CTC GAG AAT ACG TAG TGA ATA TTG GCT TTT TGA ATT-3’) was inserted into the Tet-pLKO-puro by IGEbio. The recombinant vector was cotransfected into HEK293T cells with packaging plasmids pMD2.G and psPAX2 using Lipofectamine 3000 (Life Technology, L3000015) transfection reagent. After 48 h, culture supernatant containing lentiviral particles was collected, filtered through 0.45-micron sterile filters. And then, cells were infected with lentivirus in 10 mg/mL polybrene (sc-134220, Santa Cruz Biotechnology) (52).

### EdU assay

EdU assay was performed using BeyoClick™ EdU Cell Proliferation Kit (Beyotime, C0071L). 24 h after transfection, 5000 cells were seeded in 96 well plates and cultured for another 24 h. Then, cells were incubated with EdU (20 μM) for 2 h at 37℃ and 5% CO_2_. After that, cells were fixed with 4% paraformaldehyde for 15 min, permeabilized with 0.1% Triton X-100 for 10 min. Cells were incubated with click additive solution containing Alexa fluor 488 for 30 min at room temperature in the dark, and then treated with DAPI (1:2000) for 10 min. Samples were observed and analyzed with a Lionheart FXTM Intelligent living cell imaging analysis system (BioTek, USA) (53).

### Thymidine block and Flow cytometry analysis

A 2 mg/mL thymidine (Sigma) solution was prepared in RPMI-1640 medium containing 10% fetal bovine serum. The cells were cultured with 2 mg/mL thymidine solution for 20 h, and after washing three times with PBS, cells were released with fresh culture solution for different time. Flow cytometry analysis was performed by Cell Cycle and Apoptosis Analysis Kit (Beyotime, C1052). Briefly, cells were fixed overnight in 70% ethanol at 4°C. The cells were stained with diluted propidium iodide (PI) and RNase for 30 min each at 37°C in the dark. After washing, the cells were immediately analyzed by flow cytometry using an Accuri C6 flow cytometer (5).

### DNA fiber assay

ESCC cells were transfected with siRNA for 48 h and pulse-labeled with 25 mM IdU (Sigma) for 30 min. Cells were then washed with PBS buffer twice and labeled with 250 mM CldU for 30 min. DNA fibers were spread on glass slides, and slides incubated in 2.5 M HCl for 90 min and then washed with PBS buffer. The slides were blocked in PBS containing 5% bovine serum albumin for 2 h. Primary antibodies, rat anti-BrdU antibody (Abcam, ab6326) and mouse anti-BrdU antibody (BD bioscience, 347580) were diluted in 5% bovine serum albumin and incubated for 1 h followed by extensive washing with PBS buffer. Secondary antibodies, goat anti-rat Alexa 594 and goat anti-mouse Alexa 488 were applied for 30 min and slides were mounted with antifade gold mounting media (Invitrogen). Fibers were analyzed by LSM800 with airyscan confocal laser scanning microscope (50).

### Immunoprecipitation assay

Cells were lysed on ice with EBC buffer (50 mM Tris pH 7.5, 120 mM NaCl, 0.5% NP-40, 1% protease inhibitor). Then, 2 μg of ANLN antibody (Santa Cruz, sc-271814) or normal mouse IgG (Santa Cruz, sc-2025) was added, followed by 25 μL of protein A/G magnetic beads (MedChemExpress, HY-K0202), and the mixture was incubated at 4°C for 8 h. The immunoprecipitated protein complexes were washed 5 times with EBC buffer, and the supernatant was discarded. Then 80 μL SDS loading buffer was added and boiled with 95°C for 10 min. Finally, the samples were analyzed by western blotting (54).

### Plasmids

pCMV-N-HA, pCMV-N-Flag, pcDNA3.1-3×HA, pBOBI-C-3×HA, pGEX-6P-1, pTXB1, pET-28a and pET-32a vectors were used to construct the expression plasmids. ANLN and its mutants were cloned into pBOBI-C-3×HA or pET-32a vectors. PCNA and RAD18 were cloned into pCMV-N-Flag, pCMV-N-HA, pcDNA3.1-3×HA or pGEX-6P-1 vectors. All plasmids were verified by DNA sequencing at the Beijing Genomics Institute (BGI).

### Protein purification and quantification

GST-tagged, CBD-tagged and His-tagged protein purification and quantification was performed as described (5,55). Transfect plasmid into Transetta (DE3) Chemically Competent Cell, pick positive monoclone, culture in LB medium (10 g/L tryptone, 5 g/L yeast extract, 10 g/L NaCl) at 37°C until OD_600 nm_ value of 0.7. And then, cells were induced with 0.5 mM isopropyl β-D-1-thiogalactopyranoside (Amresco, 0487) at 16°C for 16 h. After centrifugation 5,000 rpm for 5 min, the supernatant was discarded. The bacteria were lysed on ice for 30 min with GST lysis buffer (4.3 mM Na_2_HPO_4_, 1.47 mM KH_2_PO_4_, 137 mM NaCl, pH 7.3, 0.1% Triton X-100, 1 mM PMSF), CBD lysis buffer (20 mM Tris-HCl (pH 8.0), 300 mM NaCl, 0.5 mM EDTA, 1 mM PMSF) or His lysis buffer (50 mM Sodium phosphate pH 8.0, 300 mM NaCl, 10 mM Imidazole) and 1 mg/mL lysozyme (Sigma, 62971). Ultrasonic broken bacterium, centrifuge at 10,000 rpm for 10 min and collect the supernatant. In addition, the supernatant was incubated with GST Bind Resin (Millipore, 70541-5), CBD Bind Resin (Biolabs, S6651V) or His Bind Resin (Millipore, 70666-3) at 4°C for 3 h. Centrifuge at 4°C, 1, 000 rpm, and discard the supernatant. And then washed the resin 6 times with GST buffer, CBD buffer or His buffer. Finally, the resin was eluted with GST (50 mM Tris-HCl pH 8.0, and 50 mM reduced glutathione) or His eluting buffer (50 mM sodium phosphate pH 8.0, 300 mM NaCl, and 200 mM imidazole) at 4°C for 1 h; CBD Bind Resin was eluted with CBD eluting buffer (20 mM Tris-HCl (pH 8.0), 300 mM NaCl, 0.5 mM EDTA, 50 mM DTT) at 4°C for 24 h, the CBD label was removed in this process, through which we could obtain unlabeled PCNA. Centrifuge at 1000 rpm at 4 ℃ and collect the supernatant containing purified protein. All purified proteins were quantified using a Pierce BCA Protein Assay Kit (Thermo Scientific, 23225) according to the protocol provided by the manufacturer (56).

### His pulldown assay

His-tagged proteins were bound to the His-binding resin for 2 h in a His buffer (50 mM Sodium phosphate pH 8.0, 300 mM NaCl, 10 mM Imidazole) at 4°C. The resin wash with His lysis buffer three times, then pulled proteins were added and the mixture was rotated at 4°C for 4 h. Resins were washed 6 times with PBST buffer, boiled in SDS loading buffer and analyzed by western blotting (56).

### In situ proximity ligation assay

As previously described, the PLA duolink experiment was completed according to the manufacturer’s instructions (Sigma, DUO92013, DUO92005, DUO92001, DUO92005-100RXN). Cells were fixed with 4% paraformaldehyde for 15 min, permeabilized with 0.1% Triton X-100 for 10 min, then the cells were blocked with 5% donkey serum for 1 h. After that, cells were incubated overnight with antibodies against ANLN (1:200), PCNA (1:200), RAD18 (1:200) or Pol eta (1:200) overnight at 4°C. Then, the cells were incubated with diluted secondary probes at 37°C for 1 h. Next, ligation solution was added to the cells for 30 min at 37°C. After that, amplification solution was added and incubated for 100 min at 37°C. Finally, the cells were treated with DAPI (1:2000) for 10 min and observed by LSM800 with airyscan confocal laser scanning microscope (57).

### Live-cell imaging

Cells were seeded on a glass-bottom dish (Cellvis, D35-20-1.5-N). After 24 h, cells were transfected with RFP–PCNA chromobody (Chromotek, ccr, PCNA-CB) and ANLN-GFP. Then, 48h after transfection, imaging was performed at 37 °C, and 5% CO_2_ on a High Intelligent and Sensitive Microscope and every 10 min for up to 48 h (58).

### UV irradiation

The UV power of the sterilization lamp in the biosafety cabinet was quantified by using a UV measuring instrument (Linshangtech, LS126C). The cells were irradiated with different doses of UV, and the monoubiquitination level of PCNA was detected by western blotting to determine the optimal irradiation dose. UV treatment was performed when the cell density was approximately 60%-90%, and then transferred to the cell incubator for 2 h, followed by harvesting. The UV dose of 100J/m^2^ was used in this study. After testing, this dose induced the highest level of PCNA monoubiquitination in ESCC cells.

### Monoubiquitination reactions

The 4×uitination reaction buffer was prepared (80 mM HEPES (pH 7.5), 4 mM DTT, 40 mM magnesium chloride, 4 mM ATP (Roche, 10519979001)). A monoubiquitination reaction mixture containing 50 nM UBE1 (Sigma, 23-021), 50 nM Flag-RAD6B, 50 nM GST-RAD18 and 100 nM Flag-PCNA was prepared by mixing 7.5 μL of 4×ubiquitination reaction buffer with 22.5 μL of protein mixture. After incubating the reaction mixture in 30℃ for 1 h. The monoubiquitination level of PCNA was measured by western blotting (59,60).

Flag-PCNA and Flag-RAD6B was transfected into HEK293T cells, respectively. After 48h, proteins were obtained by co-immunoprecipitation and competitive elution of Flag-peptide (MCE, HY-P0223).

### Colony formation assay

The proliferation ability of ESCC cells was detected by colony formation assay. 500 cells were seeded in 1 mL complete medium on 12-well plates and incubated for 10–14 days at 37℃ with 5% CO_2_. Cells were fixed in a mixture of methanol and glacial acetic acid (3:1), stained with 0.5% crystal violet, and counted using Image J software version 1.52 (61).

### Alkaline comet electrophoresis

Spread normal melting point agar on the glass slide. The cells were diluted in PBS to 5×10^5^/mL. Take 10 μL of the cell suspension and 90 μL of 0.5% low melting point agarose and mix them at 37°C, spread on the normal melting point agar. The slides were placed in the newly prepared cell lysate (2.5 M NaCl, 100 mM Na_2_EDTA, 10 mM Tris, pH 13) at 4°C overnight. After PBS washing, the slides were placed in newly prepared alkaline electrophoresis buffer (30 mM NaOH, 2 mM Na_2_EDTA, pH 13) at room temperature for 20 min. Then the slides were electrophoresed using a horizontal electrophoresis apparatus at 25 V for 25 min. The slides were then washed with 0.4 mM Tris-HCl (pH 7.5) and stained with DAPI (1:2000) for 20 min in the dark. The sample was observed and analyzed using a fluorescence microscope (62).

### Statistical analysis

The experimental data were statistically analyzed by ImageJ and GraphPad Prism 8.0 (GraphPad Software). The mean and standard deviation (mean ± SD) of the experimental data of each group was obtained. The *t*-test was used to test the difference of the experimental data in each group (with *P* < 0.05 indicating statistical significance).

## Data availability

The data presented in this study are available in article.

## Supporting information

This article contains supporting information.

## Acknowledgements

We thank each member of the laboratory for their assistance in reagents, consumables and technical methods.

## Author contributions

Bei-Bei Tong: conceptualization, methodology, investigation, data curation, and writing-original draft. Yu-Fei Cao: formal analysis, writing-original draft, writing-review and editing. Bing-Wen: methodology and investigation. Teng-Fu: resources and software. Dan-Xia Deng: software and formal analysis. Qian-Hui Yang: Methodology. Yu-Qiu Wu: visualization. Hua-Yan Zou: resources. Lian-Di Liao: resources. Li-Yan Xu: funding acquisition, writing-review and editing. En-Min Li: funding acquisition, project administration, supervision, writing-review and editing.

## Funding and additional information

This work was supported by grants from the National Natural Science Foundation of China (81872372 and 82273108 to EML, 82173034 to LYX), the Innovative Team Grant of Guangdong Department of Education (2021KCXTD005 to EML) and 2020 Li Ka Shing Foundation Cross-Disciplinary Research Grant (2020LKSFG07B to EML).

## Abbreviations

The abbreviations used are:

ANLN: Anillin
ESCC: esophageal squamous cell carcinoma
PCNA: proliferating cell nuclear antigen
TLS: transletion synthesis
dsDNA: double-stranded DNA
RFC: Replication Factor C
polymerase: Pol
UV: ultraviolet
GSEA: gene set enrichment analysis
IdU: iododeoxyuridine
CldU: chlorodeoxyuridine
PLA: proximity alignment assay
PIP-box: PCNA-interacting proteins box
PCNA-CB: RFP–PCNA chromobody
△PIP: PIP-box deletion
CPD: cyclobutane pyrimidine dimers
Cyto: cytoplasm
Nu: nucleus
Ch: chromatin
WEC: whole-cell lysate

